# A conserved and predictable pluripotency window in callus unlocks efficient transformation in grasses and beyond

**DOI:** 10.64898/2026.01.20.700461

**Authors:** Yiyi Wang, Mengjiao Chu, Zhixia Wang, Jinhao Shao, Haijuan Zhang, Zhibiao Nan, Chunjie Li, Lei Lei

## Abstract

A major bottleneck in plant biotechnology is the inefficient and genotype-dependent regeneration of callus, which severely limits genetic transformation and functional studies across many species. This barrier is acutely exemplified in the study of beneficial plant-microbe interactions, such as the *Epichloë*-grass symbiosis—a system conferring remarkable stress tolerance to its host but hindered by a lack of efficient genetic tools. To address this, we established a chromosome-scale genome for an *Epichloë* native host grass *Achnatherum inebrians*. We discovered that the expression dynamics of evolutionarily conserved cell pluripotency regulators (CPRs) including *ARF5/7/19, BBM, WUS/WOX5* and *CUC1/2* serve as a precise molecular predictor for callus regenerative capacity, revealing that pluripotency is dynamic and peaks within a narrow, definable time window. Harnessing this predictable window enabled the development of a highly efficient transformation system for *A. inebrians* (49.4% efficiency). Crucially, this CPR-based strategy proved generalizable: applied to wheat and the legume sainfoin, it pinpointed species-specific optimal regeneration windows, boosting shoot regeneration rates to 65.7% and 87.5%, respectively. Collectively, our work provides an integrated research system and a rational design principle that removes a key barrier to uncovering molecular mechanisms in plant systems, particularly the *Epichloë*-enhanced stress tolerance symbiosis.

---

A major bottleneck in plant biotechnology is the inefficient, genotype-dependent regeneration and transformation of many species, which severely limits functional studies—particularly in agronomically important monocots. The *Epichloë*-grass symbiosis represents a prime system with high potential for enhancing crop and forage stress tolerance (Kauppinen et al. 2016). However, the elucidation of molecular mechanisms behind this endophyte-mediated tolerance has been hindered by this very constraint.

Shoot regeneration induced directly from lateral root primordia is restricted to specific developmental stages; likewise, *Arabidopsis* root explants exhibit high shoot regeneration only within a narrow window on callus induction medium (CIM) (Akama et al. 1992; Rosspopoff et al. 2017). These observations collectively indicate that callus regenerative capacity is not static, but is inherently dynamic and transient. Yet, how this capacity changes over time has not been clearly defined. Callus regeneration relies on the somatic embryogenesis to reacquire pluripotency, a process mediated by an evolutionarily conserved core transcriptional network of cell pluripotency regulators (CPRs) (Ikeuchi et al. 2019; Tang et al. 2025; Zhai and Xu et al. 2021). We therefore hypothesized that the expression dynamics of this conserved CPR network could predict the transient, critical window for regeneration.

*Achnatherum inebrians*, commonly known as drunken horse grass, naturally hosts *Epichloë* endophytes and displays tolerance to a range of biotic and abiotic stresses (Xia et al. 2018). We generated a *de novo* chromosome-scale genome assembly for *A. inebrians*, with a final size of 1.47 Gb and 60,986 annotated protein-coding genes; k-mer analysis further revealed its diploid nature and low genomic heterozygosity (∼0.47%) (Figure 1a, Figure S1). This genomic resource positions *A. inebrians* as a tractable model for studying symbiotic stress tolerance, yet realizing this goal requires an efficient transformation system. Guided by the hypothesis that CPR expression predicts regenerative capacity, we identified CPR homologs in *A. inebrians* via motif and sequence analyses of its genome, uncovering members of the ARF, PLT, and WOX families, among others (Figure S2).

**Figure 1.**
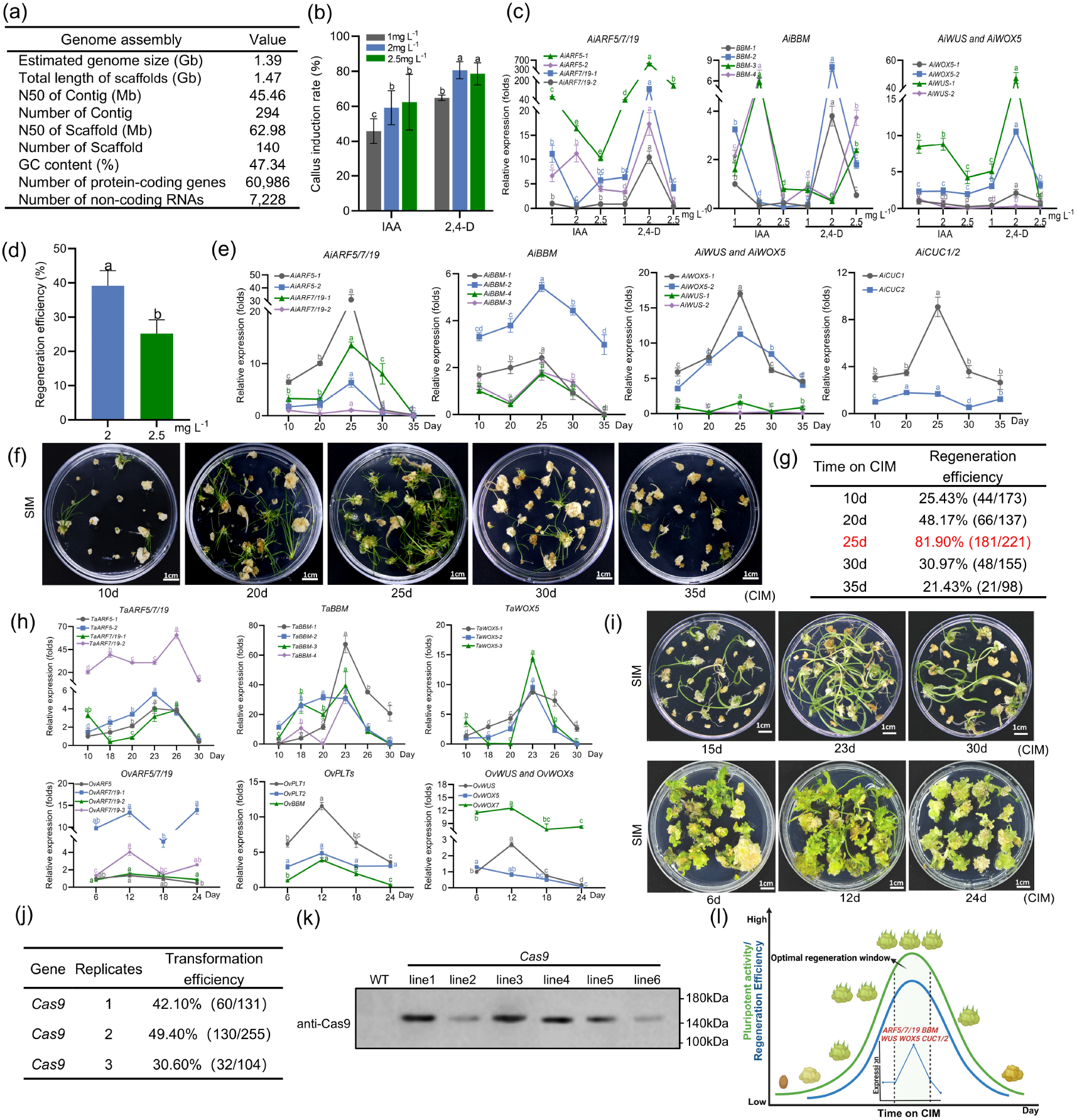
A predictable pluripotency window in callus enables high shoot regeneration and efficient transformation across species. (a) Summary of the *de novo* chromosome-scale genome assembly and annotation for *A. inebrians*. (b) Callus induction on the callus induction media (CIM) containing IAA or 2,4-D at three concentrations. (c) Expression patterns of CPR homologs (*ARF5/7/19, BBM, WUS* and *WOX5*) in *A. inebrians* calli under varying auxin concentrations. Data are means ± SD (n=3). Different letters (by color) indicate significant differences among concentrations for each gene (p < 0.05). (d) Shoot regeneration from calli after 20 d on SIM. Calli were pre-cultured on CIM supplemented with 2.0 or 2.5 mg L^−1^ 2,4-D. (e) Expression patterns of CPR homologs (*ARF5/7/19, BBM, WUS, WOX5* and *CUC1/2*) in calli over time on CIM. (f, g) Shoot regeneration capability of calli cultured on CIM for different durations. Scale bar: 1 cm. (h) Expression of conserved CPR homologs (*ARF5/7/19, BBM, WUS* and *WOX5*) in wheat and sainfoin calli over time. (i) Shoot regeneration capabilities of wheat and sainfoin calli over time. Scale bar: 1 cm. (j) *Agrobacterium*-mediated transformation efficiency in *A. inebrians*, calculated as the percentage of infected calli producing shoots. (k) Cas9 protein expression was detected in transgenic *A. inebrians* lines by western blot analysis using an anti-Cas9 antibody. (l) A proposed model predicting the optimal regeneration window based on conserved CPR expression dynamics.

To evaluate whether CPR expression levels could predict regenerative capacity, we first identified the optimal condition for embryogenic callus induction in *A. inebrians*. Three concentrations of IAA and 2,4-D were tested in CIM. Callus morphology and formation rates were similar on media containing 2 or 2.5 mg L^−1^ 2,4-D (Figure 1b, Figure S3). However, expression analysis of key CPRs revealed a clear difference; levels were high in calli induced with 2 mg L^−1^ 2,4-D but significantly lower at 2.5 mg L^−1^ (Figure 1c). This expression trend correlated with the corresponding regeneration rates (Figure 1d, Figure S3). These results demonstrate that 2 mg L^−1^ 2,4-D is optimal for induction, and confirm that CPR expression levels serve as a strong indicator of callus regenerative capacity.

To delineate the dynamic pattern of callus regenerative capacity, we profiled the expression of core CPRs in *A. inebrians* calli over a time course. The transcript levels of key regulators, including *ARF5/7/19, BBM, WUS*/*WOX5*, and *CUC1/2*, followed a concerted pattern. They increased progressively to a peak at day 25 and then underwent a sharp decline by days 30–35 (Figure 1e, Figure S4). Strikingly, this transcriptional trajectory precisely mirrored the regenerative competence of the calli (Figure 1f). Regeneration efficiency peaked (81.90%) in 25-day-old calli, then plummeted to 30.97% and 21.43% at days 30 and 35, respectively (Figure 1g). These findings demonstrate that pluripotency in callus is a transient state, defined by a narrow window of peak competence. Furthermore, they establish that the expression dynamics of evolutionarily conserved CPRs provide a predictive molecular signature for this critical regenerative window.

We further tested the generality of this principle by selecting the monocot wheat (*Triticum aestivum*) and the dicot legume sainfoin (*Onobrychis viciifolia*) for analysis. After identifying their respective CPR homologs, we monitored expression dynamics in callus. Notably, the conserved CPRs in both species exhibited a nearly identical expression trajectory that rose to a distinct peak and then declined, with peaks occurring at day 23 in wheat and day 12 in sainfoin (Figure 1h, i, Figure S5). Critically, shoot regeneration efficiency was maximized at each predicted optimum, confirming that CPR dynamics serve as a generalizable molecular predictor (Figure S5).

Leveraging the defined pluripotency-maintenance window, we developed a highly efficient *Agrobacterium*-mediated transformation platform for *A. inebrians*. Using this system, we successfully obtained transgenic seedlings expressing Cas9, which are suitable for gene editing (Li et al. 2021). This process was completed in only 45 days, with a transformation efficiency reaching 49.4% (Figure 1j, k, Figure S6).

Our work establishes that callus regenerative capacity is a transient, dynamic state, peaking within a definable window (Figure 1l). More importantly, we show that the expression dynamics of conserved CPRs serve as a predictive, species-general tool, enabling a shift from empirical screening to rational design in transformation protocol development. By integrating this predictive framework with the efficient genetic transformation system and genomic resources developed for *A. inebrians*, our study paves the way for mechanistic dissection of *Epichloë*-mediated stress tolerance in grasses.

## ACKNOWLEDGEMENTS

This study was financially supported by the National Natural Science Foundation of China (32300241, 32441036). We thank Dr. Guangpeng Ren for his advice on genome sequencing and assembly. We are grateful for Dr. Yongliang Zhang and Dr. Shicheng Yan for providing the wheat seeds.

## AUTHOR CONTRIBUTIONS

L.L., C.L. and Z.N. conceived the project. Y.W., M.C. and Z.W. performed most of the experiments and data analyses. J.S. and H.Z. provided plant material and technical support. L.L., Y.W. and M.C. wrote the manuscript. Z.N., C.L. revised the manuscript. All authors read and approved the final manuscript.

