## Supplemental Data 1 for "A conserved and predictable pluripotency window in callus unlocks efficient transformation in grasses and beyond"

### Supplemental Methods

#### Plant materials and growth conditions

The naturally *Epichloë* free of *Achnatherum inebrians* seeds were collected from Baxoi County, Xizang Autonomous Region. *A. inebrians* seeds were stratified at 4°C in darkness overnight to synchronize germination. *A. inebrians* plants were grown in a controlled environmental greenhouse at 22°C and 65% relative humidity with a 14h light/10h dark photoperiod. Wheat (Zhongmai 175) and sainfoin (Lanhong No.1) were used to test the consistency between cell pluripotency-related gene expression profiles and callus regeneration efficiency.

#### Genome size and heterozygosity estimation

Prior to assembly, the genomic characteristics of *A. inebrians* were estimated through k-mer frequency analysis of the Illumina short reads. Six-week seedlings were sampled for the DNA extraction using a CTAB method. k-mer counting was performed using jellyfish3/KMC4 based on 87.25Gb Illumina sequencing. data. The genome size was independently estimated with findGSE5 and GenomeScope6, and further corroborated by an in-house script implementing the algorithm from (Liu et al. 2013). Heterozygosity and repeat content were subsequently derived from the k-mer spectrum.

#### Genome assembly and annotation

For *de novo* genome assembly, the plant samples were prepared as described for the genome survey. The short-read sequencing was performed on the Illumina NovaSeq6000 platform with a paired-end read length of 150bp (PE150). High-fidelity long-read sequencing (HiFi) was performed on the PacBio Sequel II system. The library was constructed from high-molecular-weight genomic DNA using the SMRTbell Express Template Prep Kit 2.0 and sequenced in Circular Consensus Sequencing (CCS) mode to generate accurate reads (>99.9%), achieving over 100× coverage.

For chromosome scale genome assembly, the Hi-C library was constructed was performed following the in situ Hi-C protocol (Lieberman-Aiden et al. 2009) with minor modification. *A. inebrians* cells were cross-linked with formaldehyde. Chromatin was digested with DpnII, and the resulting sticky ends were filled in with biotinylated nucleotides. The proximate fragments were then ligated under dilute conditions. After reversing cross-links and purifying the DNA, the ligated products were sheared to 300-500 bp fragments. Fragments containing the biotinylated junctions were selectively captured using streptavidin magnetic beads. A standard Illumina sequencing library was constructed and sequenced on the NovaSeq 6000 platform to an effective depth of (>100×) genome coverage. Genome assembly was performed by integrating the Illumina short reads, PacBio HiFi long reads, and Hi-C chromatin interaction data.

Gene structure annotation was performed using an integrated approach. Transcriptomic evidence was generated from RNA-seq of leaf, stem, inflorescence, and root tissues. Predictions were derived from three complementary strategies: homology-based alignment using proteins from *Oryza sativa* and *Zea mays*; *de novo* prediction using Augustus, SNAP, and GeneMark; and transcriptome-based alignment via PASA. All evidence was integrated into a high-confidence gene set using EvidenceModeler, and its completeness was assessed with BUSCO.

**Optimization of callus induction conditions for *A. inebrians***

An adequate amount of *A. inebrians* seeds was placed in a 50 mL centrifuge tube and rinsed with 50% (v/v) sulfuric acid for 30 minutes. After discarding the acid, the seeds were washed with distilled water 5–8 times, followed by surface sterilization with 75% ethanol for 3 minutes and 4% sodium hypochlorite for 10 minutes. The seeds were then transferred to a laminar flow hood, rinsed 5–8 times with sterile water, and subjected to overnight cold stratification at 4°C. The sterilized seeds were bisected and evenly arranged on the callus induction media (CIM) containing MS (PhytoTech Labs, M519), 30 g $L^{-1}$ sucrose, 3.5 g $L^{-1}$ phytagel (Coolaber, CP8581Z),0.05 mg $L^{-1}$ 6-benzyladenine (6-BA) (Sigma, B3408), 0.6 mg $L^{-1}$ $CuSO_4$ (Sigma,C8027), and either indole-3-acetic acid (IAA) (Sigma-Aldrich, I3750) or 2,4-dichlorophenoxyacetic acid (2,4-D) (Sigma, D7299) at three concentrations (1, 2, or 2.5 mg $L^{-1}$). For each treatment, five independent biological replicates were performed. Callus induction efficiency was calculated as the percentage of seeds that produced at least one visible callus after 14 days out of the total seeds cultured per treatment.

To evaluate shoot regeneration, 14-day calli induced on CIM containing 2 or 2.5 mg $L^{-1}$ 2,4-D were transferred to shoot induction medium (SIM) consisting of MS basal medium supplemented with 3 mg $L^{-1}$ 6-benzyladenine (6-BA), 0.5 mg·$L^{-1}$ naphthaleneacetic acid (NAA) (Sigma, N640), 0.6 mg $L^{-1}$ $CuSO_4$, 30 g $L^{-1}$ sucrose, 3.5 g $L^{-1}$ phytagel. Calli were maintained at 25°C under a 16 h light/8 h dark photoperiod provided by red and yellow LEDs (70 μmol $m^{-2}$ $s^{-1}$). Shoot regeneration efficiency was determined as the percentage of calli that developed visible shoots.

**Identification of cell pluripotency-related regulator (CPR) homologs**

The candidate CPRs were selected based on their established and conserved roles in somatic embryogenesis, shoot regeneration, and the acquisition of pluripotency in model plants such as *Arabidopsis thaliana*. We focused on core transcription factors spanning multiple key pathways: auxin signaling (e.g., ARF5, ARF7, ARF19), embryonic identity specification (e.g.,LEC1, LEC2, BBM/PLT4), shoot meristem initiation and maintenance (e.g., WUS, WOX5, WOX7), lateral organ boundary establishment (e.g.,CUC1, CUC2, LBD16), and cell dedifferentiation (e.g.,WIND1, GRF4) (Ikeuchi et al. 2019; Ince and Sugimoto, 2023; Horstman et al. 2017; Zhai and Xu, 2021). These genes collectively form a conserved regulatory network whose activity is a known molecular determinant of regenerative competence.

Identification of CPR homologs in A. inebrians and other species. To identify the homologs of these candidate CPRs in A. inebrians, we performed BLASTP searches against the annotated protein database of the A. inebrians genome, using protein sequences of the above-listed Arabidopsis genes as queries. Putative homologs were further confirmed by reciprocal best-hit analysis, domain architecture verification using Pfam (ARF: PF06507 and PF02309; PLT: PF00847; WOX: PF00046), and phylogenetic analysis (see Supplementary Fig. S2 for details). The same strategy was applied to identify CPR homologs in wheat (*Triticum aestivum*) and sainfoin (*Onobrychis viciifolia*) using their respective genomic or transcriptomic databases.

**Dynamic expression analysis of CPRs**

For A. inebrians, callus samples were collected at five time points (10, 20, 25, 30, and 35 days after incubation) and immediately frozen in liquid nitrogen. Total RNA was extracted using TRIzol reagent and reverse-transcribed into first-strand cDNA. Quantitative real-time PCR (qRT-PCR) was performed using a SYBR Green-based detection system. Gene-specific primers are listed in Table S1. *AiGAPDH* was used as the internal reference gene.

Wheat callus induction was performed following previously reported protocols with minor modifications (Ishida et al. 2015; Shan et al. 2014). Callus samples were collected at seven time points (10, 15, 18, 20, 23, 26, and 30 days after incubation), immediately frozen in liquid nitrogen, and used for RNA extraction and quantitative qRT-PCR analysis. *TaACTIN* was used as the internal reference. Primers used for qRT-PCR are listed in Table S1.

*Sainfoin* seeds were surface-sterilized with 75% (v/v) ethanol for 3 min and 2% (v/v) sodium hypochlorite for 5 min, followed by air drying. Hypocotyls (5-7 cm in length) were induced at 25℃ in dark, excised into 5-mm segments, and cultured on CIM [4.43 g $L^{-1}$ MS with vitamins (PhytoTech

Labs, M519), 30 g $L^{-1}$ sucrose, 1.5 mg $L^{-1}$ 2,4-D, 0.4 mg $L^{-1}$ 6-BA, 3.5 g $L^{-1}$phytagel, pH 5.8]. Cultures were maintained at 25°C in dark, and samples were collected at 6, 12, 18, and 24 days after induction for subsequent qRT-PCR. *OvUbiquitin2* was used for normalization. Primers used for qRT-PCR are listed in Table S1.

**A. inebrians transformation system**

Surface-sterilized seeds of *A. inebrians* were bisected and cultured on CIM containing 2 mg $L^{-1}$ 2,4-D at 25°C in darkness for two weeks. Induced calli were then pre-cultured on CIM supplemented with 200 μM acetosyringone for one day. For transformation, calli were immersed in an *Agrobacterium tumefaciens* (strain EHA105) suspension ($OD_{600}$ = 0.4–0.6) for 25 minutes, blotted dry, and subsequently co-cultured on CIM with acetosyringone in darkness at 25°C for 3 days. Following co-cultivation, calli were transferred to recovery medium containing cephalosporin (500 mg $L^{-1}$) for 3–7 days. Resistant calli were selected on SIM supplemented with plasmid specific antibody. Shoot induction was conducted at 25°C under a 16-h light/8-h dark photoperiod until shoots reached 2–3 cm in length. Shoots were then transferred to root induction medium (1/2 MS salts with the same selective agent). Rooted plantlets were acclimatized *in vitro* by loosening container lids for 2 days, followed by complete removal for 3 days. Finally, agar was washed from the roots, and plantlets were transplanted into a sterilized vermiculite-soil mixture (1:1, v/v) and covered with transparent domes to maintain humidity during acclimatization.

**Wheat and sainfoin calli regeneration**

Following callus induction, the calli of wheat and sainfoin were respectively transferred onto shoot regeneration media. Wheat calli were cultured on medium containing 4.33 g $L^{-1}$ MS, 30 g $L^{-1}$ sucrose, 0.5 mg $L^{-1}$ 6-BA, 0.1 mg $L^{-1}$ NAA, and 500 mg $L^{-1}$ casein hydrolysate, solidified with 3.5 g $L^{-1}$ phytagel. Sainfoin calli were cultured on medium containing 4.43 g $L^{-1}$ MS, 30 g $L^{-1}$ sucrose, 1 mg $L^{-1}$ zeatin (Sigma, Z0164), 0.5 mg $L^{-1}$ 6-BA, solidified with 3.5 g $L^{-1}$ phytagel (pH was adjusted to 6.4). All cultures were maintained at 25°C under a 16-h light/8-h dark photoperiod. The regeneration efficiency was calculated after 20 days for wheat and after 8 weeks for sainfoin.

**Detection of Cas9 protein in transgenic *A.inebrians* plants**

To verify Cas9 protein expression, total protein was extracted from transgenic A.inebrians seedlings, with wild-type seedling serving as the negative control. Plant tissues were ground to a fine powder in liquid nitrogen and homogenized in SDS buffer (50 mM Tris-HCl pH 6.8, 1.2% SDS, 0.1 M DTT, 6% glycerol, 0.02% bromophenol blue). The homogenate was boiled for 10 min and cleaned by centrifuging at 14,000 rpm for 15 min. Protein samples were separated by SDS–PAGE and subjected to immunoblotted with a anti-Cas9 antibody (Abmart#T56852, 1:5000).

**Statistical analysis**

All experiments were performed with at least three independent biological replicates unless otherwise stated. Quantitative data are presented as mean ± standard error (SD). Statistical analyses were performed using one-way analysis of variance (ANOVA), followed by Tukey's multiple comparison test. Differences were considered statistically significant at $P < 0.05$.

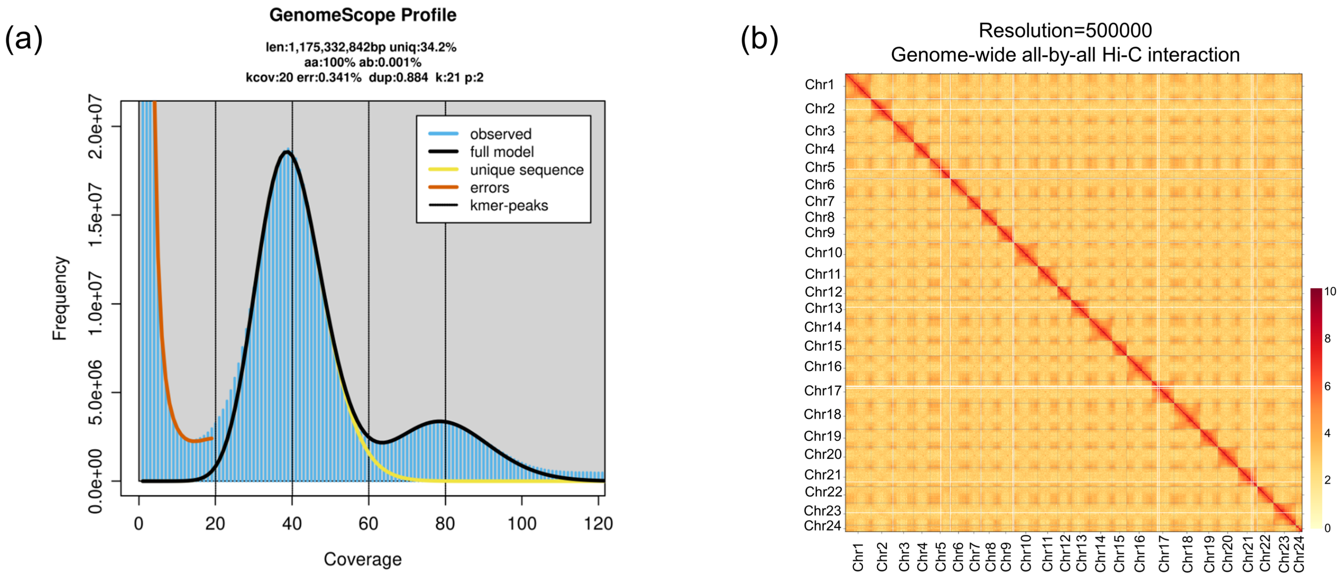


**Figure S1. Genome assembly and characterization of the *Epichloë* host grass *A. inebrians.***

**(a)** GenomeScope k-mer frequency distribution analysis of the *A. inebrians* genome. The distribution shows a heterozygous (~0.47%) genome of ~1.18 Gb.

**(b)** Hi-C interaction heatmap of the assembled *A. inebrians* genome. The plot depicts the frequency of chromatin contacts between all genomic loci at a resolution of 500Kb. The 24 assembled pseudochromosomes are labeled 1 to 24 on both axes. Interaction frequency (from low to high) is represented by the color gradient from yellow to red.

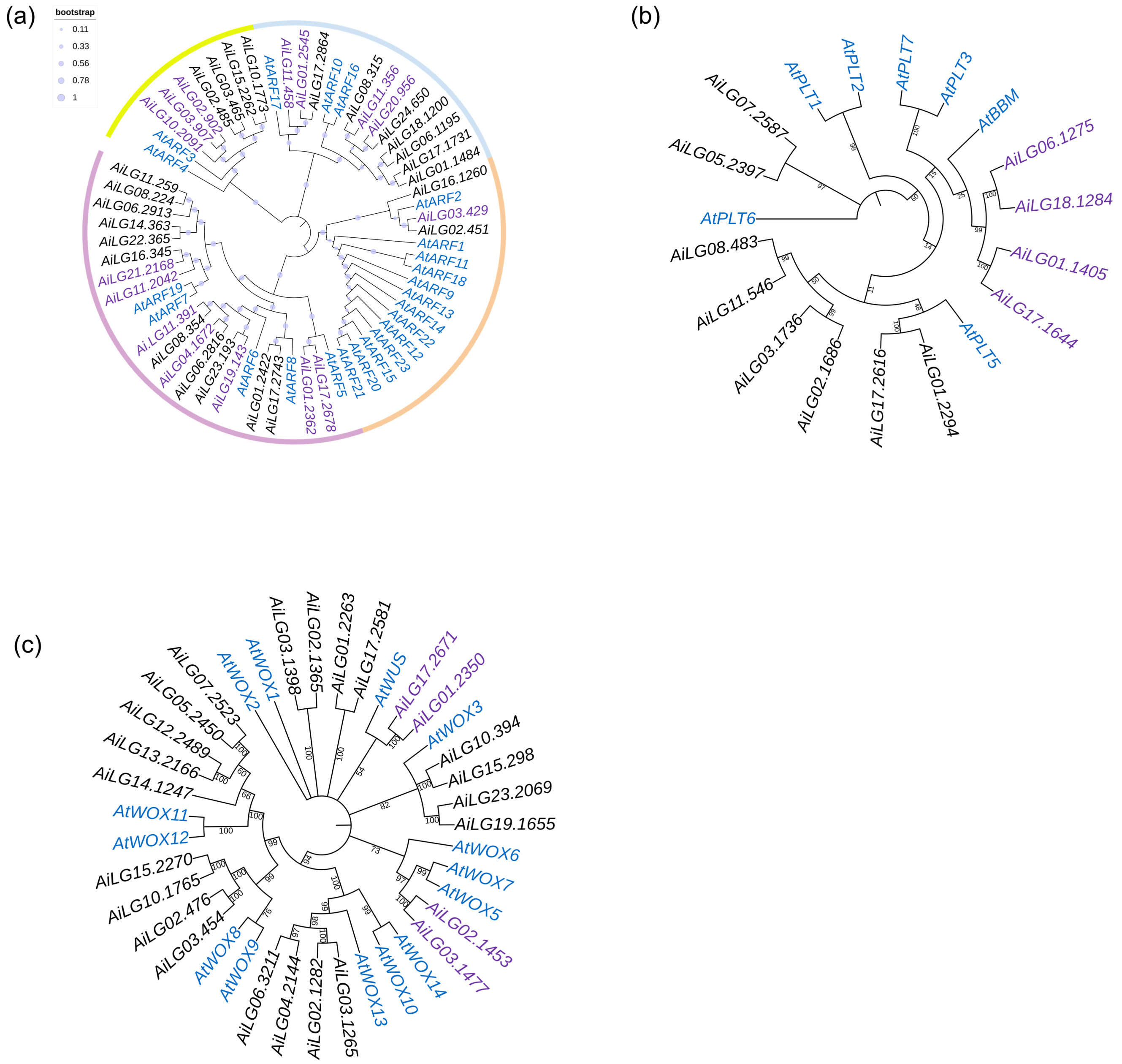


**Figure S2. Identification of *A. inebrians* ARF, PLT and WOX homologs related to core pluripotency regulators.**

**(a)** Phylogenetic analysis of the *ARF* gene families in *A. inebrians* and *Arabidopsis.*

**(b)** Phylogenetic analysis of the *PLT* gene family in *A. inebrians* and *Arabidopsis*.

**(c)** Phylogenetic analysis of the *WOX* gene family in *A. inebrians* and *Arabidopsis*. In all trees, *Arabidopsis* genes are colored blue. The closest *A. inebrians* homologs to the core CPRs are highlighted in purple.

(a)

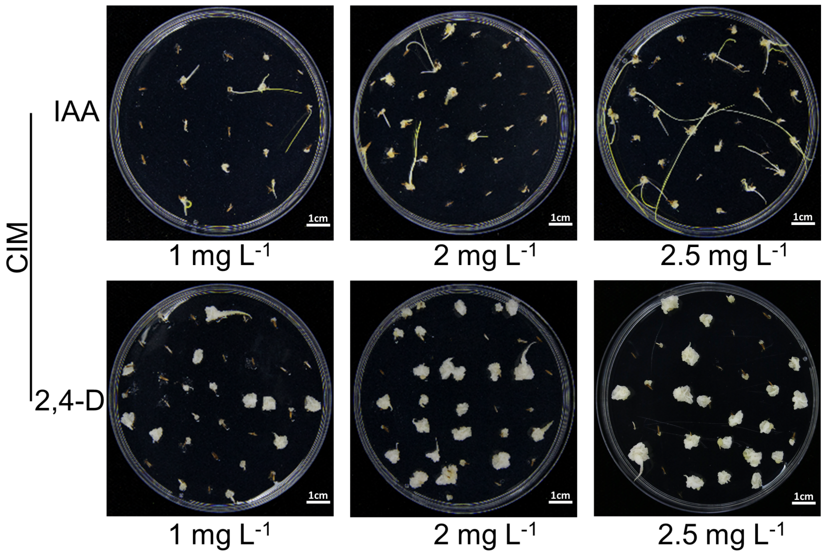


(b)

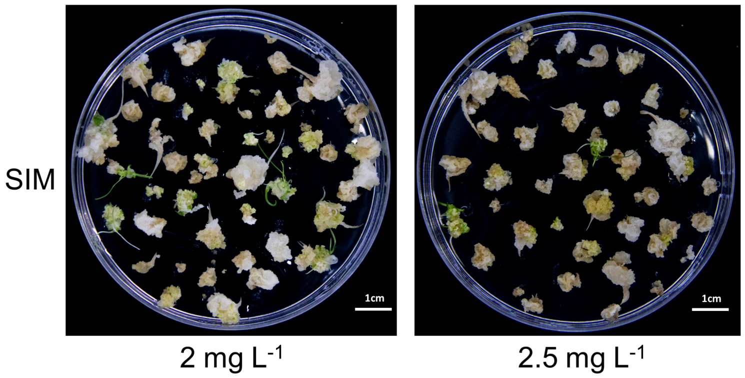


**Figure S3. Effects of different concentrations of IAA and 2,4-D in CIM on callus induction and shoot regeneration of *A. inebrians***

**(a)** Callus induction on the CIM containing IAA or 2,4-D at three concentrations. Scale bar: 1 cm.

**(b)** Shoot regeneration from calli pre-cultured on CIM with 2 or 2.5 mg $L^{-1}$ 2,4-D, after 20 days on SIM. Scale bar:1 cm.

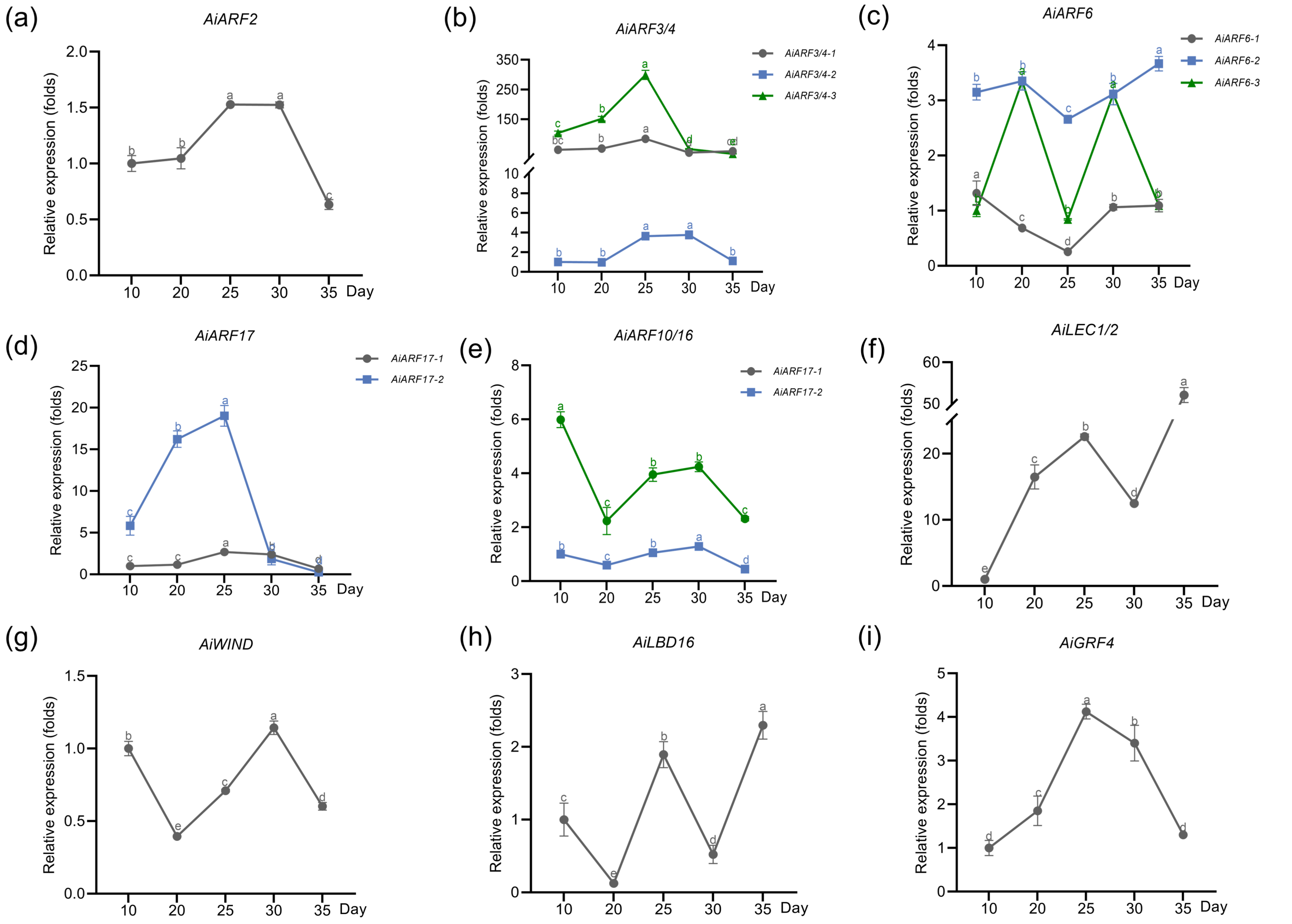


**Figure S4. Expression dynamics of *ARF* genes and CPR homologs in *A. inebrians* callus over time.**

**(a–e)** Expression patterns of *Arabidopsis ARF* homologs (excluding *ARF5/7/19*) in *A. inebrians* calli cultured on CIM at indicated time points.

**(f–i)** Expression patterns of multiple known CPR homologs under the same conditions. Gene expression was analyzed by qRT-PCR. Data represent mean ± SD ($n = 3$). For each gene, time points marked with different letters (within the same color scheme) are significantly different (one-way ANOVA with Tukey's test, $p < 0.05$).

(a)

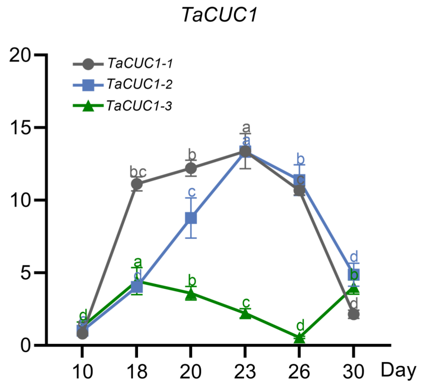


(b)

| Time on CIM | Regeneration efficiency |
|---|---|
| 10d | 40.00% (24/60) |
| 18d | 59.52% (50/84) |
| 20d | 64.63% (53/82) |
| 23d | 65.67% (44/67) |
| 26d | 65.08% (41/63) |
| 30d | 41.17% (28/68) |

(c)

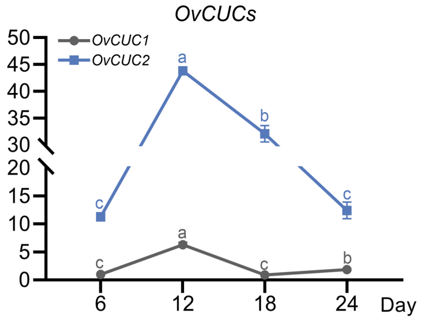


(d)

| Time on CIM | Regeneration efficiency |
|---|---|
| 6d | 70.31% (45/64) |
| 12d | 87.50% (56/64) |
| 18d | 64.06% (41/64) |
| 24d | 53.13% (34/64) |

**Figure S5. Expression of core CPR homologs predicts regenerative capacity in wheat and sainfoin callus.**

**(a)** Relative expression levels of *TaCUC1* in wheat calli cultured for different durations.

**(b)** Quantitative analysis of shoot regeneration frequency in wheat calli at corresponding time points. The condition with the highest efficiency is marked in red.

**(c)** Relative expression levels of *OvCUC1/2* in sainfoin calli cultured for different durations.

**(d)** Quantitative analysis of shoot regeneration frequency in sainfoin calli at corresponding time points. The condition with the highest efficiency is marked in red.

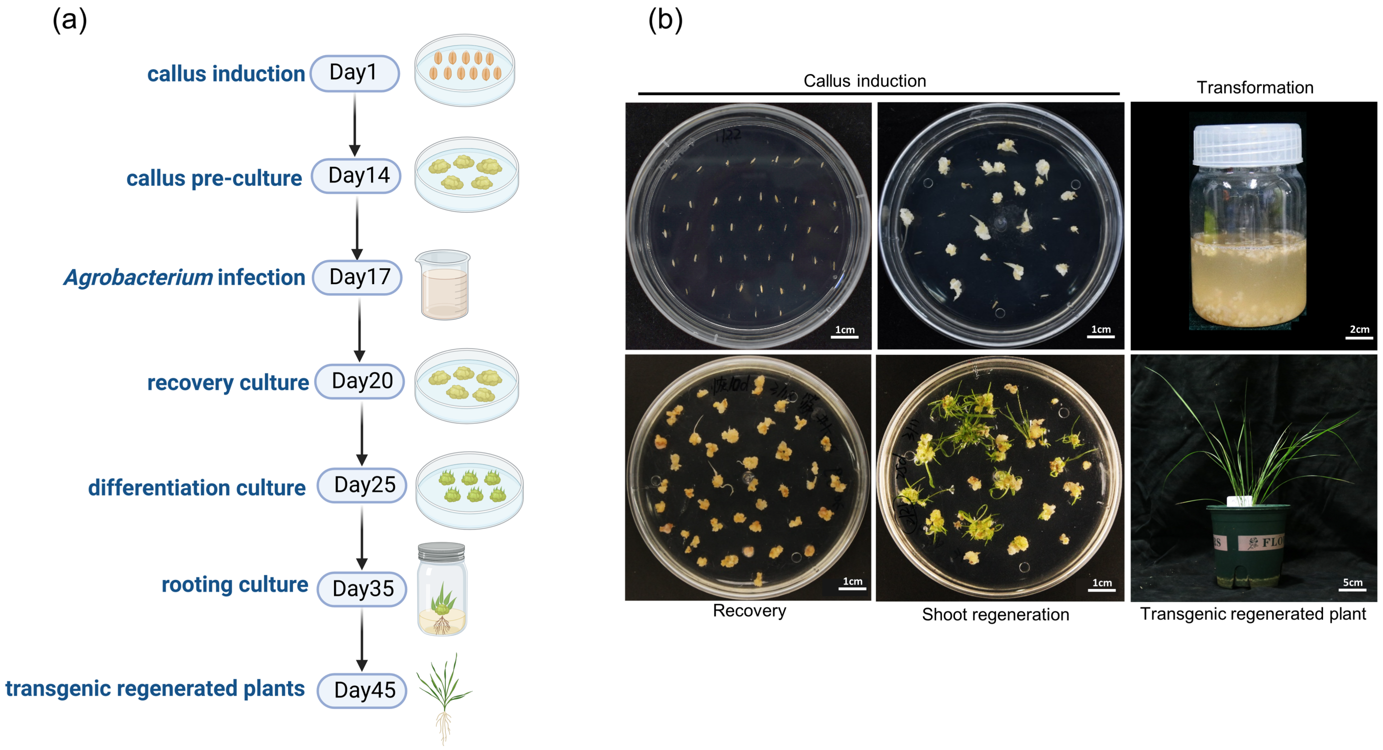


**Figure S6. Generation and characterization of Cas9-transgenic *A. inebrians* plants.**

**(a)** Flowchart depicting the key steps and time points for generating transgenic *A. inebrians* plants via *Agrobacterium*-mediated transformation.

**(b)** Representative images illustrating the key steps in generating transgenic *A. inebrians* plants: callus induction, transformation, recovery, shoot regeneration, and root regeneration. Scale bar: 1 cm; 2 cm; 5 cm.

**Table S1. Primers used in this study**

| Primer | Sequence (5’-3’) | Gene ID |
|---|---|---|
| *AiGAPDH*-F | TGTCCTTTCGTGTTCCTACTG | evm.model.LG21.1720 |
| *AiGAPDH*-R | GCAGCCTTGATAGCCTTCTT | |
| *AiARF2*-F | ATGAGGTTTGAAGGGGAAGAGG | evm.model.LG03.429 |
| *AiARF2*-R | TTGATTCGGGCCACAATGTG | |
| *AiARF3/4-1*-F | GCTTGTTGCCAAGGATTTGC | evm.model.LG02.902 |
| *AiARF3/4-1*-R | AAGATGCCTTCGTGGTTGAC | |
| *AiARF3/4-2*-F | CGGATGCTGCACATGTTCTG | evm.model.LG03.907 |
| *AiARF3/4-2*-R | TTGGGAAGGCCAAATTTCCG | |
| *AiARF3/4-3*-F | AAGCTGCGCACATTATCTGG | evm.model.LG10.2091 |
| *AiARF3/4-3*-R | CACTCCTTGGGTTCACACAAAC | |
| *AiARF5-1*-F | GCAAGCATCCCGCGG AATAC | evm.model.LG01.2362 |
| *AiARF5-1*-R | GTGTGTCCATAACATCAACCTGTGAGTTG | |
| *AiARF5-2*-F | GCAAGTGTCTTGTCTCTGTCGTCTC | evm.model.LG17.2678 |
| *AiARF5-2*-R | GTAATTGCGACTTCTCATCCCTGATGAAC | |
| *AiARF6-1*-F | GCGTTTCCGGATGCTTTTTG | evm.model.LG04.1672 |
| *AiARF6-1*-R | TGCGAGTTCTGCCAACAAAC | |
| *AiARF6-2*-F | ACAAAGCAGGTGTCATCCAC | evm.model.LG11.391 |
| *AiARF6-2*-R | GCTGCAAAAACTGACTTGGC | |
| *AiARF6-3*-F | TGGAATGCGGTTCAGAATGC | evm.model.LG19.143 |
| *AiARF6-3*-R | AACCAACCTTCACAGAACGC | |
| *AiARF7/19-1*-F | TCATTGGCGAAACCTTCAGG | evm.model.LG21.2168 |
| *AiARF7/19-1*-R | TAAAAAGGCGTGGCAACTGG | |
| *AiARF7/19-2*-F | AGTTGAAGACTGCTGCATGC | evm.model.LG11.2042 |
| *AiARF7/19-2*-R | GTTCAACAGTTGTGCTTGCG | |
| *AiARF10/16-1*-F | TTCACCTTGGAGGCT TCTTCAG | evm.model.LG11.356 |
| *AiARF10/16-1*-R | ATGGCTGGCATGTTTGACAC | |
| *AiARF10/16-2*-F | TGAAATGTGTCAGCCCATGG | evm.model.LG20.956 |
| *AiARF10/16-2*-R | TTTCGTGGCGGAGAAAATGG | |
| *AiARF17-1*-F | CGCCACCGTCATTAAGAAACAG | evm.model.LG01.2545 |
| *AiARF17-1*-R | AGGTCCAATGGTCTCAGACATC | |
| *AiARF17-2*-F | TCAAGCCAAAGTGCCCAAAG | evm.model.LG11.458 |
| *AiARF17-2*-R | TTCTGGGAGTTGGAGGCATC | |
| *AiBBM-1*-F | TACATTCGGGCAAAGGACCTC | evm.model.LG06.1275 |
| *AiBBM-1*-R | ATGAGCCTCATAACGACCTGTC | |
| *AiBBM-2*-F | TTCGTCTCCGAGCAAGATCAG | evm.model.LG18.1284 |
| *AiBBM-2*-R | AGCCAGCTCTTGATCATGGAG | |
| *AiBBM-3*-F | TTCGGACAAAGGACCTCGATC | evm.model.LG01.1405 |
| *AiBBM-3*-R | CGCCCAGATAAACTTGTTTCCC | |
| *AiBBM-4*-F | AGGCGCATCTCTGGGATAATAG | evm.model.LG17.1644 |
| *AiBBM-4*-R | CGTAACCGCCCAGATAAACTTG | |

| | | |
|---|---|---|
| *AiWOX5-1*-F | ACCACCGGTTCATGACATGC | evm.model.LG02.1453 |
| *AiWOX5-1*-R | TAGGCCTTGAGCGGGAAGAG | |
| *AiWOX5-2*-F | AGAAGCTACAACCACCACCAC | evm.model.LG03.1477 |
| *AiWOX5-2*-R | CACATCACCTCCTGCTCCAC | |
| *AiWUS-1*-F | CGTCGGAGCAGATCAGGATC | evm.model.LG17.2671 |
| *AiWUS-1*-R | ACCAGTAGAAGACGTTCTTGCC | |
| *AiWUS-2*-F | ATGCCAATGGAGGAGAAGGAAG | evm.model.LG01.2350 |
| *AiWUS-2*-R | CGCAGGATCCTGATCTGCTC | |
| *AiCUC1*-F | TTCATGCCCTACCACACCAC | evm.model.LG02.4027 |
| *AiCUC1*-R | ATCACCTTCTTGATCCCTGAGC | |
| *AiCUC2*-F | ACTCATCACGTACTACCTTCTCC | evm.model.LG08.1113 |
| *AiCUC2*-R | GGTTCGCACTTGTTCAGGTC | |
| *AiLEC1/2*-F | TACACCCCGAAAGTGAAGAGTG | evm.model.LG17.2775 |
| *AiLEC1/2*-R | CAACAACCGGGGTTTGATCTAG | |
| *AiLBD16*-F | CCACATCTTCGCCCTCCAG | evm.model.LG07.3609 |
| *AiLBD16*-R | AGCTGCATCATCCCTTGCC | |
| *AiGRF5*-F | TACGCGTACTATGGGAAGAAGC | evm.model.LG11.2515 |
| *AiGRF5*-R | AGGCTTTCTTGAACGGTTGC | |
| *AiWIND*-F | ACATCAACACTCCATCCTACGC | evm.model.LG06.535 |
| *AiWIND*-R | AAGAGTACAGTGGAGGAAGCTG | |
| *TaActin*-F | GCCGTTCTGTCCTTGTATGC | AY145451 |
| *TaActin*-R | CCTGACCATCAGGCATCTCA | |
| *TaARF5-1*-F | TCAAGCAAGCGCAGATACAC | TraesCS2A02G491000 |
| *TaARF5-1*-R | AGTTGCGCCATTTGGAGTTG | |
| *TaARF5-2*-F | AGCAAAATGGCAAGGGTGAG | TraesCS2B02G519200 |
| *TaARF5-2*-R | TGTTGGTCATCGCAAAACCG | |
| *TaARF7/19-1*-F | AATGGCGCAACATCAACAGC | TraesCS7B02G065800 |
| *TaARF7/19-1*-R | AATTGGGCTTGCTGTTGCTG | |
| *TaARF7/19-2*-F | TGGCTCAACATCAACAGCAG | TraesCS7D02G161900 |
| *TaARF7/19-2*-R | AACCGCAGAACTTTGCTGTG | |
| *TaWOX5-1*-F | TCTGCAGCTTCGATCGGTATC | TraesCS3A02G368100 |
| *TaWOX5-1*-R | ACAAGCAGCTACATGCATGC | |
| *TaWOX5-2*-F | ATCCCCGGATCATGACATGC | TraesCS3B02G399800 |
| *TaWOX5-2*-R | AGGTCGTAGGCTGCTTTGAG | |
| *TaWOX5-3*-F | ACAGGTTCAGCCGGTATGTG | TraesCS3D02G361100 |
| *TaWOX5-3*-R | ATACCGATCGAAGCTGCAGAG | |
| *TaBBM-1*-F | ACATATGACCAGGCAGGAGTAC | TraesCS2B02G378100 |
| *TaBBM-1*-R | TGATGCGCCACGAGAAAAAC | |
| *TaBBM-2*-F | TGACTAGGCAGGAATACATCGC | TraesCS6A02G229500 |
| *TaBBM-2*-R | CAACCCTCCCAATTCTTGCTTG | |
| *TaBBM-3*-F | GGTTGCAGGAAACAAGGATCTC | TraesCS6B02G252000 |
| *TaBBM-3*-R | AAGTTGGTGACGGCATTGAG | |
| *TaBBM-4*-F | ATCAGCCTTCATACGGGATCAC | TraesCS6D02G205300 |

| | | |
|---|---|---|
| *TaBBM-4*-R | TCACCCCTCATGTTCCAATCG | |
| *TaCUC1-1*-F | GCATGAAGAAGACGCTCGTC | TraesCS7A02G334800 |
| *TaCUC1-1*-R | TTGTTGGGCAGGAAGTGGTAG | |
| *TaCUC1-2*-F | AACCTCAGTCTGCCGATGTTC | TraesCS7B02G246300 |
| *TaCUC1-2*-R | TGCCCTCCAGCTTAATGTCG | |
| *TaCUC1-3*-F | AGTGGGTCATGCACGAGTTC | TraesCS7D02G342300 |
| *TaCUC1-3*-R | CTCATCCCTTGTGGTGTTGTTG | |
| *OvUbiquitin 2*-F | GCTACGGTTTGGACGCAGACCATAC | PB.13954.1 |
| *OvUbiquitin 2*-R | CCTCCTCCTCCCATCCCATTTGTG | |
| *OvARF5*-F | GAGACAGAGGAATCCGGTAAAC | PB.13163.1 |
| *OvARF5*-R | CAGGAGTCTCAATCTCCCAAAC | |
| *OvARF7/19-1-F* | CTTCAGGTTGGATGGGATGAA | PB.22573.1 |
| *OvARF7/19-1*-R | CTTCATGCCAAGGTCATCTCC | |
| *OvARF7/19-2*-F | GAATCCACAGCCGGTGAGAG | PB.822.1 |
| *OvARF7/19-2*-R | GCCCTCGATCCGCATCTCG | |
| *OvARF7/19-3*-F | GTGCGGCGATACATGGGTAC | PB.4900.1 |
| *OvARF7/19-3*-R | GGTAATCCCGGAGGCCATGG | |
| *OvPLT1*-F | GGCTCTCAGCTTGGACAGCATC | PB.1847.1 |
| *OvPLT1*-R | GTTCTTCTCCTCCCATTTGAGGC | |
| *OvPLT2*-F | GAGGGGTGACAAGGCACCAC | PB.6828.2 |
| *OvPLT2*-R | GCTTCAGCTGCTTCCTCTTG | |
| *OvBBM*-F | GTTCTTCTCCTCCCATTTGAGGC | PB.4184.1 |
| *OvBBM*-R | GGTCCCTGCAGATTCACCTTC | |
| *OvWUS*-F | GATCCACCCTCACCACAGGG | PB.12608.1 |
| *OvWUS*-R | CCGTACTTGCTAAGCTGAACAG | |
| *OvWOX5*-F | CTGGCTCCTCCTCTGACTCTG | PB.25444.1 |
| *OvWOX5*-R | CCGTACTTGCCAAGCTGAACAG | |
| *OvWOX7*-F | CCGTACTTGCTAAGCTGAACAG | PB.15499.1 |
| *OvWOX7*-R | GGACCGTGCTGACTCAAGTC | |
| *OvCUC1*-F | GGGAGAGCACCAAAGGGAG | PB.24936.1 |
| *OvCUC1*-R | GGATCCACAAGAGGAGGGAGTG | |
| *OvCUC2*-F | GGAAAGCCCCCAAAAGGTG | PB.11758.2 |
| *OvCUC2*-R | CCACTGTTTCTGCCTCTGTTG | |
